# PC1-guided transcriptomic stratification reveals hepatic transcriptional heterogeneity and defines a myeloid-associated 20-gene signature

**DOI:** 10.64898/2026.08.14.744782

**Authors:** Zihan Li, Fuheng Xie, Yuqing He, Li Ma, Qiang Liu

## Abstract

Hepatic lipid-associated inflammation contributes to metabolic liver disease and cardiometabolic complications. Treatment-based transcriptomic comparisons can obscure inter-individual heterogeneity when animals exposed to the same experimental condition show divergent molecular responses. In the public hyperlipidemic liver transcriptomic dataset GSE338111, conventional sex-adjusted comparison of Amlexanox versus DMSO identified only 21 differentially expressed genes at FDR < 0.05 and |log₂FC| ≥ 1, and submission of this DEG set to Metascape yielded no GO Biological Process enrichment result. We therefore applied treatment-independent, PC1-guided transcriptomic stratification based on the 500 most variable genes. This analysis resolved three PC1-derived groups and enabled derivation of a myeloid-associated 20-gene signature from the G2-versus-G1 contrast. Independent bulk-transcriptomic cohorts supported responsiveness of the signature to dietary challenge and pharmacologic intervention, while single-cell analysis localized its expression predominantly to hepatic myeloid populations. Human cis-eQTL Mendelian randomization and colocalization further identified TAGLN2 as the signature gene with the strongest genetic support for coronary heart disease. Together, these findings show that PC1-guided stratification can improve resolution of heterogeneous hepatic transcriptional responses and provide a cross-cohort molecular signature for subsequent mechanistic and translational evaluation.

## Introduction

Hepatic lipid dysfunction is a central feature of metabolic dysfunction-associated steatotic liver disease (MASLD), the current consensus nomenclature for the condition formerly termed NAFLD [1,2]. MASLD is highly prevalent worldwide and is closely linked to type 2 diabetes, metabolic syndrome, and cardiovascular disease, including coronary artery disease [3–7,9]. Disease progression reflects interacting disturbances in lipid handling, insulin action, hepatocellular stress, and intrahepatic immune activation, with hepatic macrophages emerging as important regulators of inflammatory and repair responses [8,10–12]. Murine hyperlipidemic and steatotic liver models remain major preclinical platforms for studying lipid-associated hepatic injury and evaluating candidate interventions, although model-specific features can influence translational interpretation [13–16]. RNA sequencing and single-cell approaches now enable biological heterogeneity to be resolved at transcriptome-wide and cellular levels, while dimensionality reduction and cross-dataset integration provide complementary tools for defining reproducible molecular states [17,19,21,22].

Nevertheless, treatment-based grouping remains common in hepatic metabolic transcriptomic studies. PCA is widely used for dimensionality reduction and visualization [17,19], but pronounced inter-individual structure within nominal groups can complicate treatment-based comparisons. When substantial molecular heterogeneity exists within nominal experimental groups, limited replication and heterogeneous responses can reduce differential-expression power and compromise the reproducibility of molecular signatures [20,21]. Sex can also contribute substantially to transcriptomic variation and therefore warrants explicit consideration in mixed-sex molecular studies [23]. In the GSE338111 cohort, which was generated to examine sex-specific effects of Amlexanox in a PCSK9/Paigen-diet model [25], conventional analysis identified only 21 stringent DEGs, and Metascape analysis of this DEG set yielded no GO-BP enrichment result. These observations motivated a treatment-independent examination of the dominant transcriptomic structure before subsequent subgroup analysis. More broadly, external validation and explicit assessment of heterogeneity and potential confounding are important for improving the reproducibility and translational interpretation of molecular signatures [21,26].

To address these analytical challenges, we developed a multi-layer workflow centered on PC1-guided transcriptomic stratification. Rather than assuming that treatment allocation fully captured the dominant molecular structure, the workflow first examined treatment-independent variation across the cohort and then derived a compact 20-gene signature from the resulting PC1-derived groups. The signature was subsequently evaluated in independent dietary, pharmacologic, and single-cell datasets and was further examined using human genetic analyses. This framework is intended to complement conventional treatment-based analysis by improving the resolution of heterogeneous hepatic transcriptional responses.

## Methods

### 1. General experimental design

This study performed a retrospective bioinformatics reanalysis of public RNA-seq datasets using a unified workflow. Briefly, the analysis comprised: (1) conventional differential expression based on the original treatment grouping; (2) treatment-independent PCA and PC1-guided transcriptomic stratification; (3) screening of a 20-gene signature using predefined statistical thresholds; (4) external evaluation in independent bulk and single-cell transcriptomic datasets; and (5) human genetic analysis to assess translational relevance. Key filtering thresholds and multiple-testing procedures were prespecified within each analytical module, and analysis code was retained to support reproducibility.

### 2. Dataset acquisition and preprocessing of GSE338111

The bulk RNA-seq dataset GSE338111 was retrieved from NCBI GEO. In the original study, hyperlipidemic stress was induced in C57BL/6 mice with AAV8-mediated hepatic Pcsk9 overexpression [25]. The dataset contained five vehicle-treated (DMSO) and six Amlexanox-treated mice. The deposited gene-level count-like values contained non-integer entries; these values were rounded to non-negative integers only for DESeq2-based analyses. Lowly expressed genes were removed by requiring a count ≥ 10 in at least 3 samples. Sex was included as a covariate in the primary differential-expression models.

### 3. Differential expression analysis under original treatment grouping

Differential expression between Amlexanox and vehicle groups was analyzed using DESeq2 [18] with sex included as a covariate. DEGs were defined as FDR < 0.05 and |log₂FC| ≥ 1. A volcano plot summarized genome-wide transcriptional changes. The resulting stringent DEG set was submitted to Metascape [53] for GO Biological Process enrichment analysis.

### 4. PCA-based sample stratification and identification of the 20-gene signature

Variance-stabilizing transformation (VST; blind = TRUE) was applied to the filtered count matrix using DESeq2 [18], followed by PCA using the 500 genes with the greatest variance across all 11 mice. Based on their distribution along PC1, samples were assigned to three PC1-derived groups: G1 (n = 3), G2 (n = 4), and G3 (n = 4). G1 contained one DMSO-treated and two Amlexanox-treated mice, G2 contained four DMSO-treated mice, and G3 contained four Amlexanox-treated mice. All subsequent subgroup analyses used these same PC1-derived assignments.

A sex-adjusted DESeq2 comparison between G2 and G1 [18] was used for signature screening. Genes with FDR < 0.05 and |log₂FC| ≥ 3 were retained, ranked by mean normalized expression (baseMean), and the top 20 were defined as the core 20-gene signature. VST expression values of the signature genes were converted to gene-wise Z-scores for heatmap visualization under the original treatment and PC1-derived grouping schemes. Based on the resulting signature profiles, G1 was designated Low-signature, G2 as High-signature, and G3 as Intermediate-signature.

For sample-level scoring, the mean normalized expression of each signature gene in G2 was used as the reference, with a pseudocount of 0.5 added to numerator and denominator. The final score was calculated as the log₂-transformed geometric mean of gene-wise relative expression ratios. Two-sided Wilcoxon rank-sum tests were used for group comparisons.

Comparisons among the PC1-derived groups were adjusted using the Benjamini–Hochberg procedure; the original DMSO-versus-Amlexanox comparison was treated as an exploratory single comparison.

### 5. Transcriptome quality control, sex-adjustment sensitivity, and Pcsk9-related transcriptional auditing

Quality-control assessment included library size, the number of detected genes, and sample-wise VST expression distributions. Pairwise Spearman correlation matrices were displayed using the same transcriptome-wide VST expression matrix, with samples ordered first by the original treatment assignment and then by the PC1-derived groups. Sensitivity to sex adjustment was assessed by comparing genome-wide log₂FC estimates from group-only and sex-adjusted DESeq2 models [18] for the G2-versus-G1 and G3-versus-G2 contrasts.

As an independent transcriptomic audit outside the 20-gene panel, log₂-transformed Pcsk9 normalized counts (pseudocount = 1) and three non-overlapping modules representing PCSK9/cholesterol homeostasis, FXR/bile acid signaling, and lipogenesis were examined across the PC1-derived groups. Module scores were summarized from direction-adjusted gene-wise Z-scores. A descriptive comparison restricted to female DMSO-treated samples was additionally performed without inferential testing because of the small and unbalanced sample numbers.

### 6. Differential expression analysis across PCA-defined subgroups

Pairwise DESeq2 comparisons [18] (G2 versus G1 and G3 versus G2) used FDR < 0.05 and |log₂FC| ≥ 1 to define DEGs. Volcano plots highlighted all 20 signature genes. GO Biological Process over-representation analysis was performed with clusterProfiler [52] using the significant DEGs from each contrast and all tested Entrez-mappable genes as the background, followed by Benjamini–Hochberg correction and semantic redundancy reduction using Wang similarity (cutoff = 0.50). An effect-size heatmap summarized the two contrasts for the 20-gene signature, and six representative genes were displayed using log₂-transformed normalized counts (pseudocount = 1) as individual-mouse trajectories.

### 7. Single-cell localization of the 20-gene signature (GSE235939)

The mouse liver single-cell dataset GSE235939 [41] was used to characterize the cell-type distribution of the 20-gene signature. Cells were retained when detected genes ≥ 200, total RNA counts were 250–5000, and mitochondrial transcripts comprised < 5% of total counts. Processing was performed with Seurat v5, and sample-level integration followed the established Seurat integration framework [22]. After log normalization, 3000 highly variable genes were used for PCA and graph-based clustering, with sample-level integration used for the common UMAP representation. Broad cell identities were manually assigned by examining canonical lineage-marker expression together with cluster-specific marker genes identified using Seurat FindAllMarkers. Conservative broad labels were used where finer annotation was uncertain. Gene-expression visualization used the RNA assay rather than the integrated assay. Dot plots and UMAP feature plots summarized cell-type-specific expression. No individual cell was treated as an independent biological replicate for bulk-style hypothesis testing.

### 8. Dietary cohort validation (GSE287727)

GSE287727 comprised WT and HuRKO mice fed chow or a high-fat, high-cholesterol, high-fructose (HFCF) diet (n = 3 per group). Genes were retained when counts were ≥ 10 in at least 3 samples. A factorial DESeq2 model [18] (∼ genotype + diet + genotype:diet) was used to estimate dietary effects in WT and HuRKO mice. Where multiple Ensembl identifiers mapped to the same candidate-gene symbol, the identifier with the highest mean normalized expression was retained. Sample-level signature scores were calculated as the mean of the available gene-wise Z-scores. Genotype, diet, and interaction effects were evaluated by linear modeling and ANOVA; planned HFCF-versus-Chow contrasts within each genotype were obtained with the R package emmeans and adjusted across the two diet comparisons using the Holm method. Gene-level forest plots displayed DESeq2 log₂FC estimates with Wald 95% confidence intervals; FDR < 0.05 was used for gene-level significance, with ±1 log₂FC shown as an effect-size reference.

### 9. Pharmacological cohort validation (GSE235797)

The GSE235797 dataset [41] included 3 vehicle and 3 Ginkgetin-treated high-fat diet mice. Duplicate gene entries were averaged, and zero-variance genes were removed before analysis.

Limma [51] with empirical Bayes moderation was used for differential expression, with Benjamini–Hochberg FDR correction. Sample-level signature scores were compared using a two-sided Welch t-test. Volcano plots, clustered heatmaps, and forest plots summarized genome-wide and signature-level responses. This cohort was used as an independent directional pharmacologic validation rather than as a direct head-to-head comparison with Amlexanox.

### 10. Human Mendelian randomization and colocalization analysis

Mouse signature genes were mapped to human one-to-one orthologs; Ccl6 was excluded from the primary analysis because no direct human ortholog was available. Genetic instruments were derived from GTEx v8 liver and whole-blood cis-eQTL data [24], with cis variants defined within ±1 Mb of the transcription start site of the corresponding gene. Variants were filtered at P < 5×10⁻⁸ and F-statistic > 10 and were LD-clumped at r² < 0.001 within 10,000 kb. Alleles were harmonized between exposure and outcome summary statistics. Wald ratio estimates were used for single-instrument analyses and inverse-variance weighted estimates when multiple instruments were available [42,43]. Disease outcome summary statistics were obtained from FinnGen NAFLD (894 cases and 217,898 controls; N = 218,792; European ancestry), GCST90091033 NAFLD (8,434 cases and 770,180 controls; N = 778,614; European ancestry), and CARDIoGRAMplusC4D coronary heart disease (60,801 cases and 123,504 controls; N = 184,305; mixed ancestry, predominantly European). The historical NAFLD labels of the source GWAS were retained when referring to these datasets.

Bayesian colocalization was performed using eQTLGen whole-blood data [46] within ±500 kb of the lead eQTL, with regions containing fewer than 50 shared SNPs excluded.

Colocalization inference followed the coloc framework [44]. PP.H4 ≥ 0.80 was considered strong evidence and 0.50 ≤ PP.H4 < 0.80 moderate evidence for a shared causal variant.

Benjamini–Hochberg correction was applied to MR results within the prespecified outcome families.

### 11. Statistical analysis and software

All analyses were performed in R version 4.6.1. Bulk differential expression analyses used DESeq2 [18] or limma [51], single-cell processing used Seurat [22], functional enrichment used clusterProfiler [52], and MR/colocalization analyses used TwoSampleMR/MR-Base [43] and coloc [44]. Benjamini–Hochberg correction was applied within analytical modules where multiple testing was performed. Unless otherwise specified, statistical tests were two-sided.

FDR < 0.05 was considered statistically significant and unadjusted P < 0.05 nominally significant. Individual mice were treated as biological replicates in bulk-transcriptomic comparisons, and 95% confidence intervals were reported where applicable. Fixed random seeds were used for stochastic analyses.

## Results

### 1. Transcriptomic profiling of Amlexanox intervention under hyperlipidemia

Conventional treatment-based differential analysis of GSE338111 was first performed using the prespecified thresholds of FDR < 0.05 and |log₂FC| ≥ 1. The sex-adjusted Amlexanox-versus-DMSO comparison identified only 21 DEGs, including 18 upregulated and 3 downregulated genes (Supplementary Fig. S1A). When these 21 DEGs were submitted to Metascape, no GO Biological Process enrichment result was returned, providing limited functional resolution under the original treatment allocation. Together with the substantial overlap of DMSO- and Amlexanox-treated animals in PCA space, this limited differential signal motivated treatment-independent examination of inter-individual expression heterogeneity.

### 2. PC1-guided stratification identifies distinct expression states and establishes the 20-gene signature

Unsupervised PCA based on the 500 most variable genes was performed across all 11 GSE338111 samples. PC1 and PC2 explained 64.8% and 21.4% of the total transcriptional variance, respectively. Under the original treatment labels, DMSO- and Amlexanox-treated mice overlapped substantially in PCA space (Fig. 1A). In contrast, the samples occupied three visually distinct regions along PC1, while PC2 visually separated male and female samples (Fig. 1B). According to the PC1 distribution, the rightmost region was denoted G1 (n = 3), the leftmost region G2 (n = 4), and the intermediate region G3 (n = 4). G1 comprised one DMSO-treated and two Amlexanox-treated mice, whereas G2 and G3 comprised four DMSO-treated and four Amlexanox-treated mice, respectively. The analysis workflow is summarized in Fig. 1C. A sex-adjusted G2-versus-G1 comparison was then used for signature derivation: genes with FDR < 0.05 and |log₂FC| ≥ 3 were retained, ranked by baseMean, and the top 20 were selected. The resulting expression profiles characterized G1 as Low-signature, G2 as High-signature, and G3 as Intermediate-signature. Heatmaps showed the lowest overall expression in G1, the highest expression in G2, and an intermediate pattern in G3 (Fig. 1D). The sample-level signature score showed the same ordering, whereas the original DMSO-versus-Amlexanox comparison was not significant (P = 0.055; Fig. 1E). These findings indicate that PC1-guided stratification provided greater transcriptomic resolution than the original binary treatment grouping.

**Figure 1.**
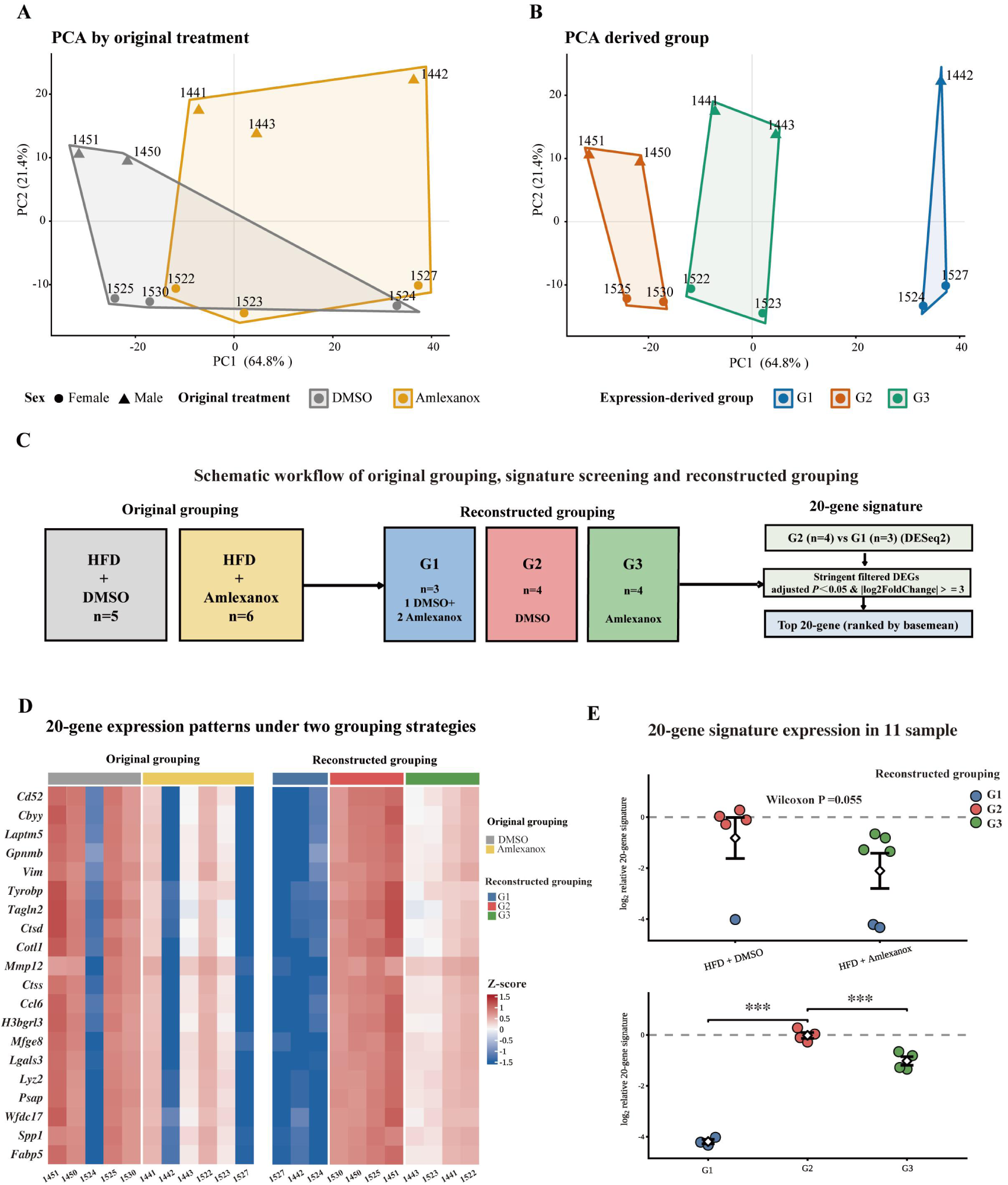
Identification of a 20-gene transcriptional signature and reconstruction of PC1-derived subgroups in GSE338111. (A) Principal component analysis (PCA) of the 11 GSE338111 liver transcriptomes colored by the original treatment assignment. PCA was calculated after blind variance-stabilizing transformation using the 500 genes with the greatest across-sample variance. Sex is indicated by point shape. PC1 and PC2 explain 64.8% and 21.4% of the total variance, respectively. (B) The same PCA coordinates displayed according to the three PC1-derived groups, denoted G1 (n = 3), G2 (n = 4), and G3 (n = 4). (C) Schematic overview of the original treatment grouping, PC1-guided reconstruction of the three groups, and subsequent derivation of the 20-gene signature. The cohort comprised HFD-fed mice treated with DMSO (n = 5) or Amlexanox (n = 6). G1 contained one DMSO-treated and two Amlexanox-treated mice, G2 contained four DMSO-treated mice, and G3 contained four Amlexanox-treated mice. After PC1-guided grouping, candidate genes were identified by sex-adjusted DESeq2 analysis of G2 versus G1, filtered at FDR < 0.05 and |log2 fold change| ≥ 3, ranked by mean normalized expression (baseMean), and the top 20 genes were selected. Subsequent signature profiling characterized G1 as Low-signature, G2 as High-signature, and G3 as Intermediate-signature. (D) Heatmaps of the same 20-gene expression matrix displayed according to the original treatment grouping (left) and the PC1-derived grouping (right). Expression values are row-wise Z scores of variance-stabilized expression. (E) Comparison of the 20-gene signature under the original treatment grouping (top) and PC1-derived grouping (bottom). Each point represents one mouse and is colored according to the PC1-derived group; white diamonds indicate group means and error bars indicate SEM. The signature represents the log2-transformed relative expression score derived from the 20-gene panel. The original DMSO-versus-Amlexanox comparison was assessed using a Wilcoxon rank-sum test. Comparisons among G1, G2, and G3 were assessed using pairwise Wilcoxon rank-sum tests with Benjamini-Hochberg correction. Asterisks denote adjusted significance levels: *adjusted P < 0.05, **adjusted P < 0.01, and ***adjusted P < 0.001.

### 3. Transcriptome quality control, sex-adjustment sensitivity, and Pcsk9-related transcriptional auditing

Quality-control metrics showed broadly comparable detected-gene numbers and VST expression distributions across GSE338111 samples despite variation in library size (Supplementary Fig. S1B). Pairwise Spearman correlation matrices, shown under the original treatment ordering and the PC1-derived group ordering, further illustrated the cohort-wide similarity structure (Supplementary Fig. S1C). Genome-wide effect estimates were highly concordant between group-only and sex-adjusted models for both subgroup contrasts, supporting the robustness of the principal differential-expression effects to inclusion of sex as a covariate (Supplementary Fig. S1D). Bulk Pcsk9 levels were comparable among G1, G2, and G3, indicating no significant difference in upstream Pcsk9-driven hyperlipidemic stimulation intensity and transgenic overexpression efficiency across subgroups. Notably, G1 exhibited uniform and stable Pcsk9 transcription, whereas G2 and G3 showed prominent intra-group transcriptional fluctuation. Gene expression-based functional module scoring revealed unchanged transcriptional levels of upstream cholesterol-homeostasis pathways across subgroups, but significantly activated downstream bile acid signaling and lipogenesis pathways at the transcriptomic level in G1 (Supplementary Fig. S1E-G). These results rule out inconsistent upstream hyperlipidemic stimulation as the driver of subgroup divergence and reveal distinct downstream transcriptional profiles in bile acid signaling and lipogenesis among the PC1-derived hepatic groups.

### 4. Subgroup-specific transcriptional shifts during expression-state transition and Amlexanox treatment

Sex-adjusted pairwise differential analysis (FDR < 0.05, |log₂FC| ≥ 1) characterized subgroup-specific transcriptional changes. The G2-versus-G1 comparison yielded 4,609 DEGs (2,524 higher and 2,085 lower in G2) (Fig. 2A). These DEGs were enriched for GO Biological Processes related to leukocyte activation, cytokine production, phagocytosis, leukocyte adhesion, and migration, placing the 20-gene signature within a broader immune and myeloid-associated transcriptional context (Fig. 2B). The G3-versus-G2 comparison, corresponding to Amlexanox versus DMSO treatment within these PC1-derived groups, identified 1,262 DEGs (863 higher and 399 lower in G3) (Fig. 2C). All 20 signature genes were reduced in G3 versus G2, of which 12 were significantly downregulated. GO analysis of the G3-versus-G2 DEGs identified enrichment of regulation of tumor necrosis factor production, myeloid leukocyte activation, and myeloid leukocyte differentiation (Fig. 2D).

**Figure 2.**
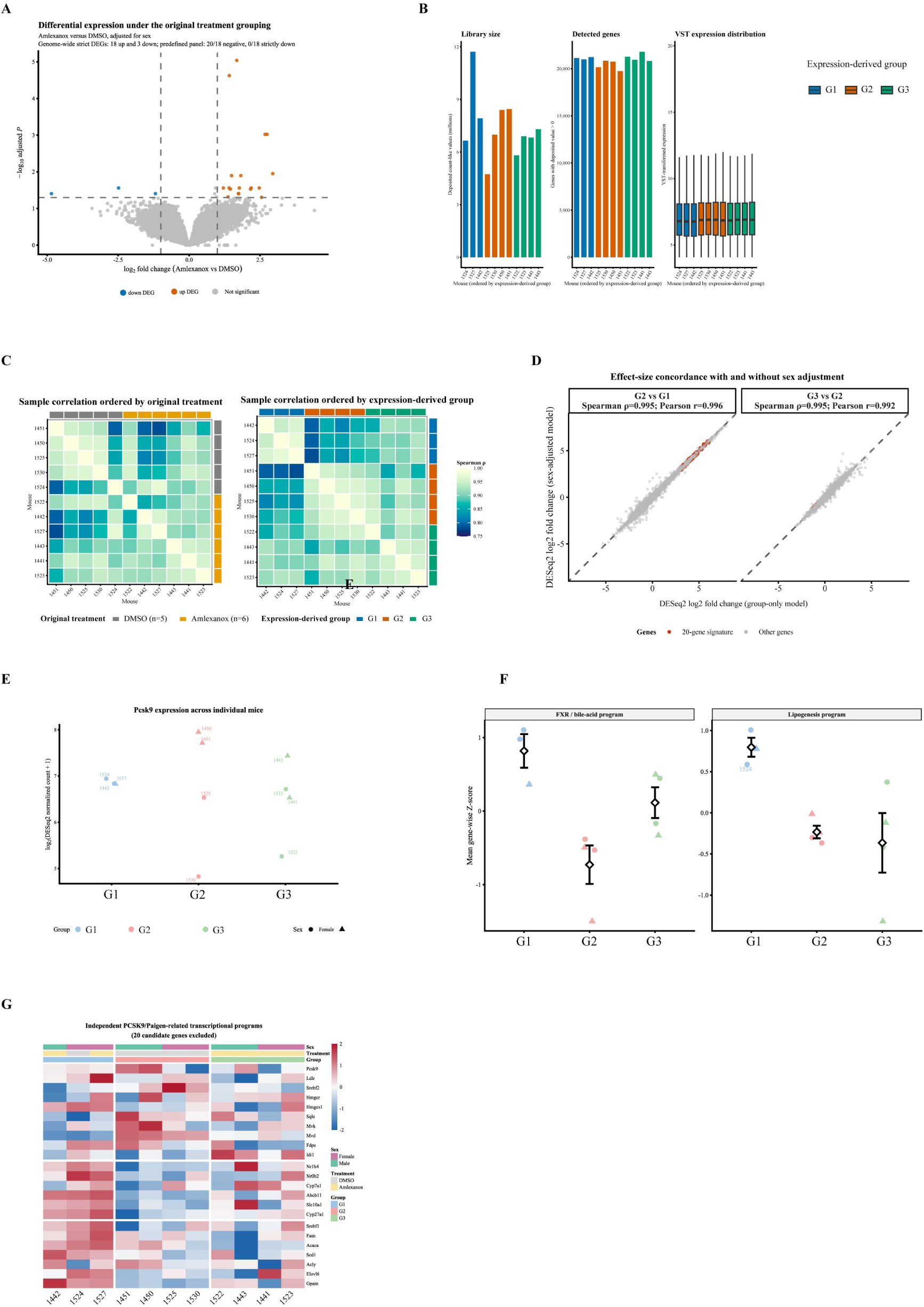
Transcriptomic characteristics of the PC1-derived groups of GSE338111. (A) Volcano plot of differential gene expression between G2 (High-signature) and G1 (Low-signature). (B) Gene Ontology Biological Process enrichment analysis of significant differentially expressed genes between G2 and G1. (C) Volcano plot of differential gene expression between G3 (Intermediate-signature) and G2 (High-signature); G3 comprised Amlexanox-treated mice and G2 comprised DMSO-treated mice. (D) Gene Ontology Biological Process enrichment analysis of significant differentially expressed genes between G3 and G2. For volcano plots, differentially expressed genes were defined as FDR < 0.05 and |log2 fold change| ≥ 1; genes belonging to the predefined 20-gene signature are highlighted and labeled. Dashed vertical lines indicate log2 fold changes of -1 and +1, and the horizontal dashed line indicates FDR = 0.05. GO analyses included all significant up- and downregulated genes from the corresponding comparison; bubble size represents gene count and color represents -log10(FDR). (E) Effect-size heatmap of the 20-gene signature showing DESeq2 log2 fold changes for G2 versus G1 and G3 versus G2. Red and blue indicate positive and negative log2 fold changes, respectively. Asterisks indicate FDR significance: *FDR < 0.05, **FDR < 0.01, and ***FDR < 0.001. (F) Expression trajectories of six representative signature genes (Mfge8, Cybb, Sh3bgrl3, Ctsd, Gpnmb, and Ccl6) across G1, G2, and G3. Each point represents one mouse; gray lines connect group means, white diamonds indicate means, and error bars indicate SEM. Expression is shown as log2(normalized count + 1). Differential-expression analyses were performed using DESeq2 with sex included as a covariate.

Effect-size and trajectory plots further showed strong G1-to-G2 induction followed by a consistent G2-to-G3 decrease across representative signature genes (Fig. 2E, F). Thus, PC1-guided stratification revealed a biologically coherent transcriptional contrast that was less apparent under the original treatment grouping.

### 5. Single-cell transcriptomics localizes the signature predominantly to hepatic myeloid populations

Independent single-cell RNA-seq data from GSE235939 [41] were used to characterize the cellular distribution of the 20-gene signature. Among the profiled non-parenchymal liver cell populations, most signature genes were preferentially expressed in myeloid subsets, particularly Kupffer/macrophage and inflammatory monocyte/macrophage populations (Fig. 3A). Individual genes also showed broader or distinct distributions across other immune and stromal populations. UMAP visualization resolved the major annotated non-parenchymal cell populations (Fig. 3B), and feature plots for Cybb, Ccl6, Tagln2, and Wfdc17 further illustrated the predominantly myeloid-associated pattern (Fig. 3C). Because GSE235939 mainly characterizes hepatic non-parenchymal cells, the observed enrichment supports predominantly myeloid localization of this 20-gene signature within the profiled non-parenchymal compartment. The small residual hepatocyte population showed low detection proportions and weak expression of the signature genes.

**Figure 3.**
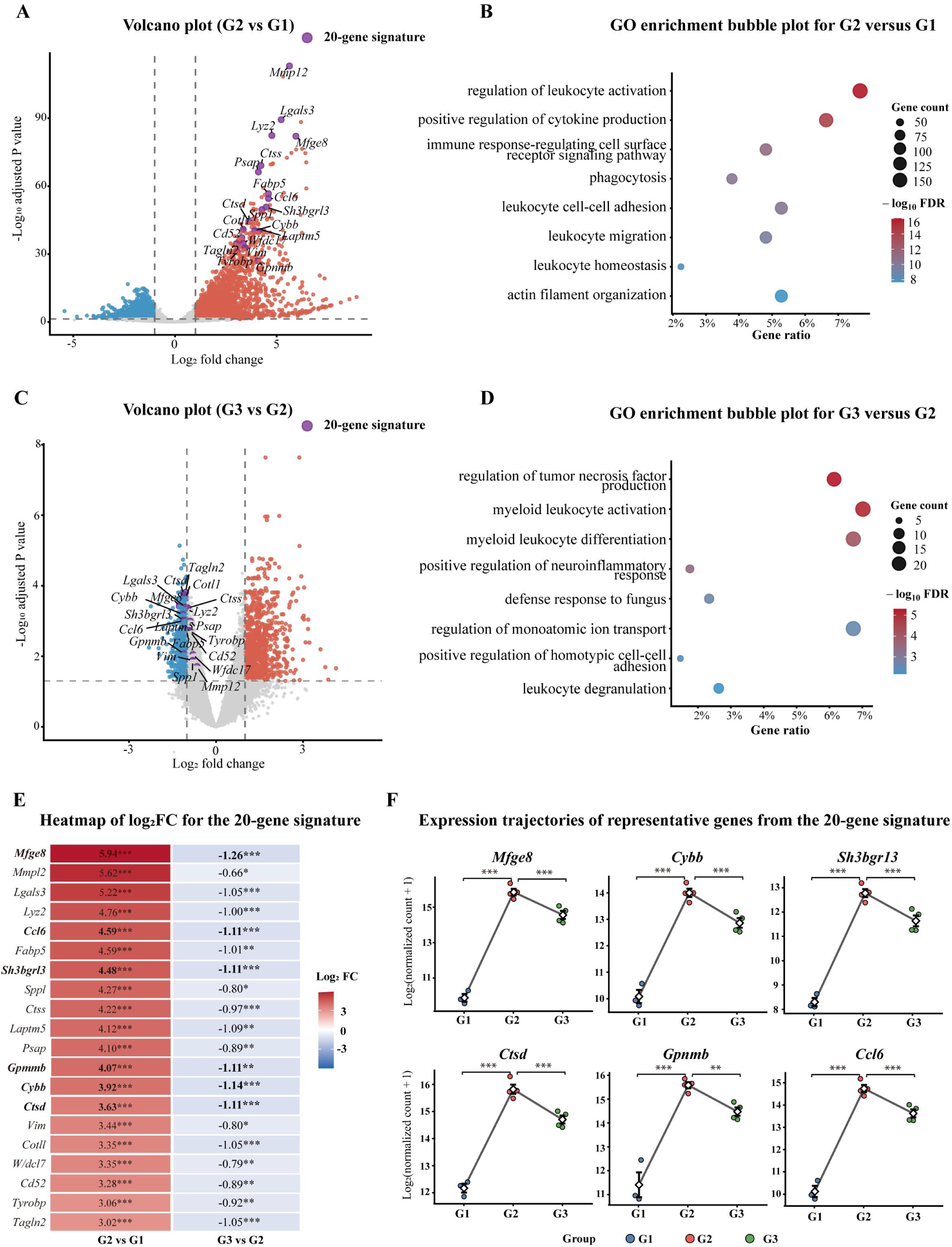
Single-cell localization of the 20-gene transcriptional signature in GSE235939. (A) Dot plot showing the expression distribution of the 20-gene signature across major liver cell populations identified in the single-cell RNA-sequencing dataset. Dot size represents the percentage of cells expressing each gene, whereas color represents scaled average expression within each cell type. (B) UMAP representation of the single-cell transcriptomes annotated into broad cell populations, including Kupffer/macrophages, inflammatory monocytes/macrophages, DC-like myeloid cells, pDCs, endothelial cells, stellate/mesenchymal cells, cholangiocytes, T cells, B cells, plasma cells, NK cells, cycling cells, and hepatocyte-contaminating cells. Sample-level reciprocal PCA integration was used exclusively for visualization of the UMAP embedding. (C) UMAP feature plots showing normalized RNA expression of four representative signature genes (Cybb, Ccl6, Tagln2, and Wfdc17) across the full single-cell atlas. Cells with no detectable expression are shown in light gray; expression intensity among positive cells is represented by the color gradient. For each gene, the upper visualization limit was set to the 97th percentile of positive expression values to reduce the influence of extreme values. Expression-based analyses in panels A and C were performed using the original RNA assay rather than the integrated expression matrix.

### 6. Independent dietary cohort supports signature responsiveness to HFCF challenge

The dietary cohort GSE287727, comprising WT and HuRKO mice fed chow or HFCF diets, was used for external evaluation of the 20-gene signature. HFCF feeding increased signature scores in both genotypes, with a larger overall shift in HuRKO mice (Fig. 4A). The heatmap showed broad diet-associated induction of the signature, although one WT-HFCF mouse displayed a Low-signature-like pattern resembling G1 in the primary cohort, consistent with an attenuated individual transcriptional response to HFCF (Fig. 4B). Gene-level effect estimates were predominantly positive for HFCF versus Chow, with particularly large effects for Gpnmb and Mmp12; Fabp5 showed the most discordant direction among the panel genes (Fig. 4C). These results provide independent support for responsiveness of the signature to dietary lipid stress while also illustrating inter-individual heterogeneity within the external cohort.

**Figure 4.**
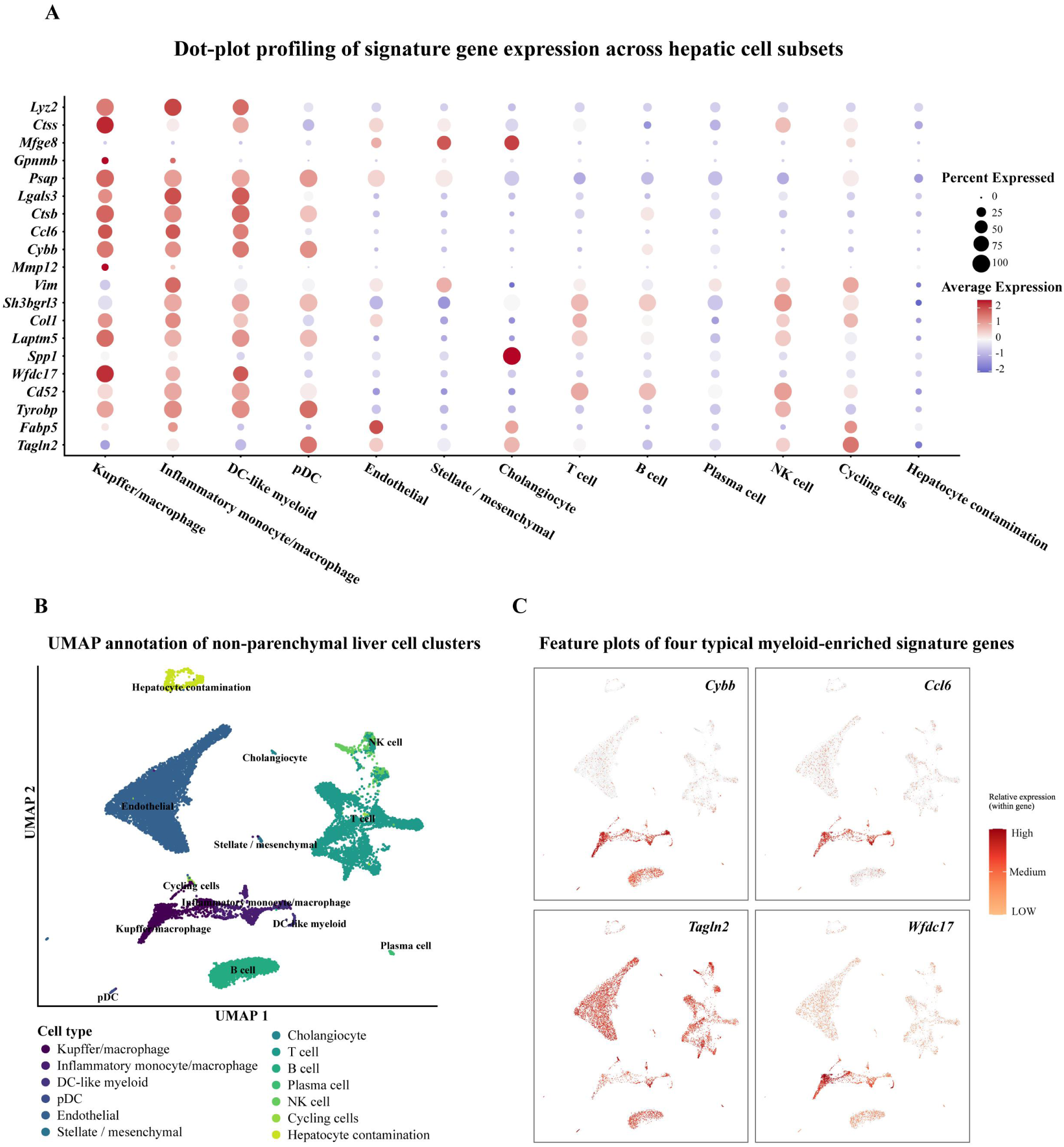
External validation of the 20-gene transcriptional signature in the independent GSE287727 MASLD model. (A) Twenty-gene signature scores across WT-Chow, WT-HFCF, HuRKO-Chow, and HuRKO-HFCF mice. Signature scores were calculated as the mean gene-wise Z score of the available 20-gene panel. Each point represents one mouse, white diamonds indicate group means, and error bars indicate SEM. Effects of genotype, diet, and genotype × diet interaction were evaluated by linear modeling and ANOVA; planned HFCF-versus-Chow comparisons within genotype were adjusted across the two diet contrasts using the Holm method. (B) Heatmap showing gene-wise Z-scored expression of the 20-gene signature across the four diet/genotype groups. Columns are ordered as WT-Chow, WT-HFCF, HuRKO-Chow, and HuRKO-HFCF to maintain adjacent comparisons within genotype; genes were hierarchically clustered. Red and blue indicate relatively higher and lower expression, respectively. (C) Gene-level diet-associated effect estimates for the 20-gene panel within WT and HuRKO mice. The two panels show WT-HFCF versus WT-Chow and HuRKO-HFCF versus HuRKO-Chow, respectively. Points represent DESeq2 log2 fold-change estimates and horizontal lines indicate Wald 95% confidence intervals. Positive values indicate higher expression under HFCF feeding and negative values indicate lower expression. Point colors distinguish FDR-significant from nonsignificant increases and decreases, with FDR < 0.05 considered significant. DESeq2 analyses used a genotype × diet factorial model.

### 7. Independent pharmacologic cohort supports signature suppression after Ginkgetin treatment

The Ginkgetin intervention cohort GSE235797 [41] was analyzed as an independent pharmacologic validation dataset. Ginkgetin significantly reduced the sample-level 20-gene signature score (Welch t-test, P = 0.0026; Fig. 5A). Genome-wide limma analysis and the signature heatmap showed a coordinated treatment-associated shift (Fig. 5B, C). All 20 signature genes had negative effect estimates after treatment, and 18 remained significantly downregulated after FDR correction. Forest-plot estimates confirmed broad suppression across the panel, with Mmp12 and Gpnmb among the genes showing the largest decreases (Fig. 5D). Because Ginkgetin and Amlexanox were evaluated in separate experimental systems, these data provide cross-treatment directional support for pharmacologic suppression of the signature rather than a direct comparison of drug efficacy.

**Figure 5.**
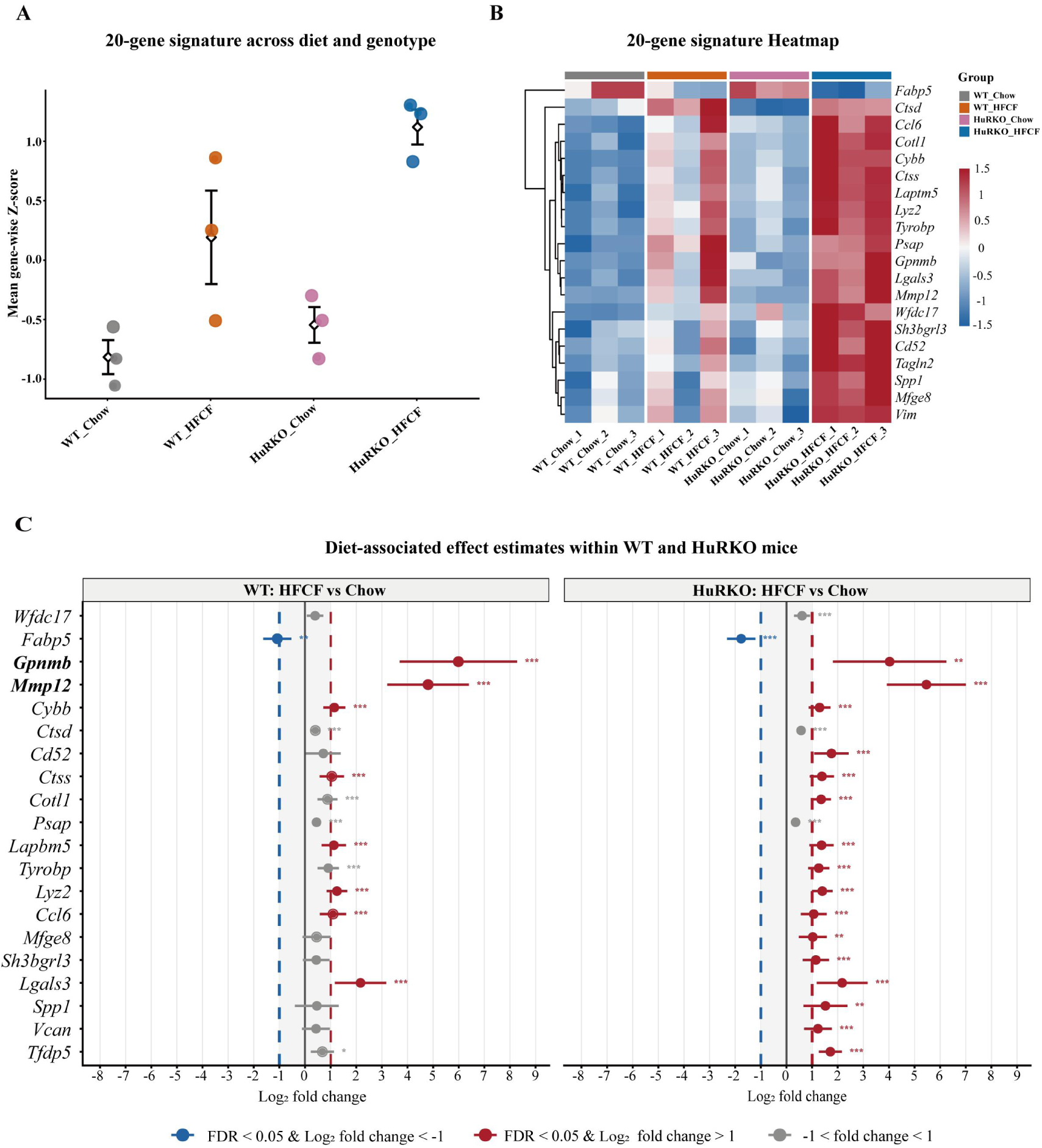
External pharmacologic validation of the 20-gene transcriptional signature in GSE235797. (A) Twenty-gene signature scores in Vehicle- and Ginkgetin-treated mice. Signature scores were calculated as the mean gene-wise Z score across candidate genes detected in the external dataset. Each point represents one mouse; white diamonds indicate group means and error bars indicate SEM. Groups were compared using Welch’s t-test (P = 0.0026). (B) Volcano plot of Ginkgetin-associated differential expression determined by limma. Candidate genes from the 20-gene signature that were detected in GSE235797 are highlighted and labeled. Other differentially expressed genes were defined as FDR < 0.05 and |log2 fold change| ≥ 1. Dashed vertical lines indicate log2 fold changes of -1 and +1, and the horizontal dashed line indicates FDR = 0.05. (C) Heatmap showing gene-wise Z-scored expression of the available 20-gene signature genes across Vehicle- and Ginkgetin-treated mice. Red and blue indicate relatively higher and lower expression, respectively. (D) Gene-level treatment effect estimates for the available signature genes. Points indicate limma log2 fold-change estimates for Ginkgetin versus Vehicle, and horizontal lines indicate moderated 95% confidence intervals. Negative values indicate lower expression following Ginkgetin treatment. Point colors distinguish FDR-significant and nonsignificant upward or downward effects. Differential-expression analysis was performed using limma with empirical Bayes moderation and Benjamini-Hochberg adjustment for multiple testing.

### 8. Human genetic analysis identifies TAGLN2 as a translational candidate

To explore translational relevance, mouse signature genes were mapped to human orthologs, with CCL6 excluded from the primary analysis because no direct human ortholog was available. GTEx v8 liver and whole-blood cis-eQTLs were used as genetic instruments for Mendelian randomization (MR). Five gene-tissue pairs showed nominal associations (P < 0.05), and whole-blood TAGLN2 was the only signal that remained significant after FDR correction (Fig. 6A, B). The two source fatty-liver GWAS were historically labeled NAFLD, whereas MASLD is used elsewhere in this manuscript for current disease nomenclature.

**Figure 6.**
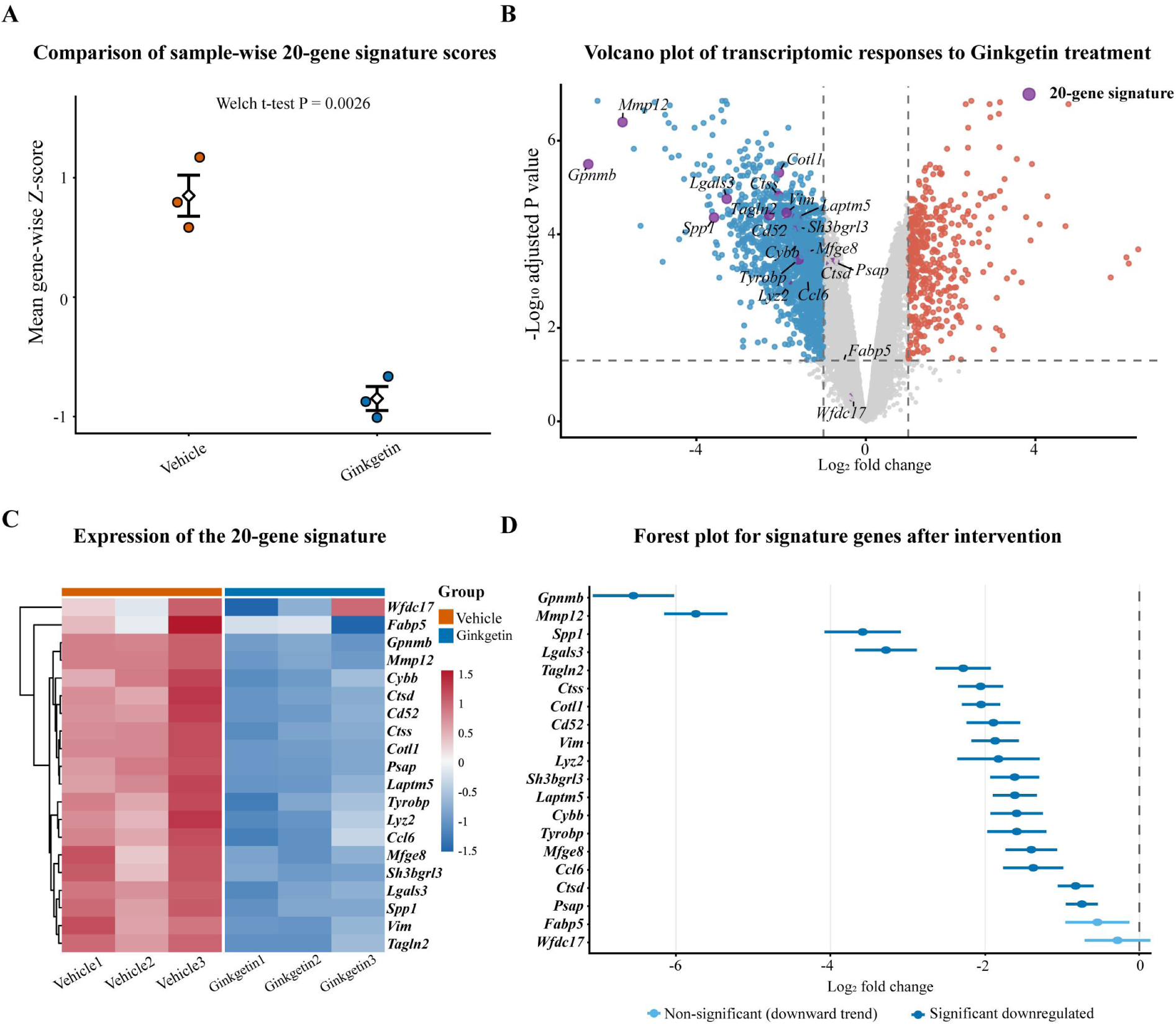
Human genetic analysis of the 20-gene signature by Mendelian randomization and colocalization. (A) Mendelian randomization screening of candidate human orthologs across FinnGen NAFLD, GWAS Catalog NAFLD, and coronary heart disease outcomes. The x-axis shows log2(odds ratio) and the y-axis shows -log10(P). Circles indicate liver eQTL instruments and triangles indicate whole-blood eQTL instruments. Point colors distinguish FDR-significant associations (FDR < 0.05), nominal associations (P < 0.05 and FDR ≥ 0.05), and nonsignificant associations. The vertical dashed line denotes OR = 1 and the horizontal dashed line denotes P = 0.05. (B) Heatmap summarizing MR evidence across gene-tissue-outcome combinations. Red and blue indicate log2(OR) values above and below zero, respectively; ** denotes FDR < 0.05 and * denotes nominal P < 0.05. (C) Forest plot of coronary heart disease MR estimates. Points represent odds ratios per genetically predicted increase in gene expression and horizontal lines indicate 95% confidence intervals; point shape indicates eQTL tissue and point color indicates statistical evidence as in panel A. The vertical dashed line denotes OR = 1. (D) Regional colocalization analysis of TAGLN2 whole-blood expression and coronary heart disease. Regional eQTLGen and GWAS association signals are shown across the shared genomic interval, with rs2789422 highlighted as the SNP with the highest SNP-level posterior support (SNP PP.H4 = 1.000). The accompanying posterior-probability plot summarizes hypotheses H0-H4; PP.H4 = 0.9966 indicates strong evidence consistent with a shared causal variant.

Across the NAFLD outcomes, the other signature genes showed weaker and less consistent MR evidence. For coronary heart disease, TAGLN2 showed the strongest human genetic association among the tested gene-tissue pairs (Fig. 6C). Regional colocalization at the TAGLN2-CHD locus yielded PP.H4 = 0.9966, providing strong evidence consistent with a shared causal variant; rs2789422 carried the highest SNP-level posterior support under the shared-variant hypothesis (Fig. 6D). These results prioritize TAGLN2 as the signature gene with the strongest combined MR and colocalization support for CHD, without establishing direct causal regulation by gene expression alone.

## Discussion

Treatment-based transcriptomic comparisons can lose resolution when substantial inter-individual molecular heterogeneity exists within nominal experimental groups, particularly in small RNA-seq cohorts where biological variation has a strong influence on differential-expression power [17,19–21]. In GSE338111, the conventional sex-adjusted DMSO-versus-Amlexanox comparison yielded only 21 stringent DEGs, and Metascape analysis of this DEG set returned no GO-BP enrichment result; meanwhile, the two treatment groups overlapped extensively in PCA space. These observations do not imply that treatment-based analysis is intrinsically invalid; rather, they show that treatment allocation alone did not capture the dominant transcriptomic structure of this cohort. PC1-guided stratification therefore provided a complementary way to resolve that heterogeneity and enabled identification of a coherent 20-gene expression program. This interpretation is consistent with broader recommendations to examine biological heterogeneity explicitly and to test molecular signatures across independent settings rather than relying on a single discovery contrast [21,26].

To clarify the potential drivers of divergent transcriptional phenotypes across uniformly hyperlipidemic mice, we analyzed Pcsk9 expression and core hepatic pathway activities at the transcriptomic level. Bulk Pcsk9 expression and upstream cholesterol-homeostasis signaling were comparable among G1, G2, and G3, indicating equivalent transgenic overexpression efficiency and consistent upstream hyperlipidemic stimulation across all samples. Notably, G1 exhibited stable and uniform Pcsk9 transcription, whereas G2 and G3 displayed prominent intra-group transcriptional fluctuation. Downstream functional scoring revealed significantly activated bile acid signaling and lipogenesis pathways in G1, while upstream cholesterol homeostasis remained unchanged. These findings exclude variable external lipid stress as the cause of subgroup divergence and support intrinsic downstream transcriptional reprogramming as a major contributor to the observed hepatic heterogeneity. AAV-mediated expression of gain-of-function PCSK9 is an established approach for inducing sustained hypercholesterolemia in wild-type mice, including in atherosclerosis-oriented models [16], and the source GSE338111 study used this framework to characterize sex-dependent hepatic responses to Amlexanox [25]. The broader metabolic actions of Amlexanox through inhibition of IKKepsilon/TBK1 have also been demonstrated in experimental models and in a clinical metabolic study [27,28], while lipid handling, bile-acid signaling, insulin resistance, and inflammatory responses are recognized as tightly interconnected components of steatotic liver disease biology [8–10].

Single-cell mapping further showed that the 20-gene signature was predominantly represented in hepatic myeloid populations within the profiled non-parenchymal compartment. This finding is consistent with the central role and marked functional heterogeneity of Kupffer cells and recruited monocyte-derived macrophages in metabolic liver disease [11,12,29,39].

Single-cell atlases have also demonstrated substantial macrophage and broader cellular heterogeneity in the human liver [36,37]. Disease-focused single-cell and spatial studies have identified macrophage states associated with TREM2, CD9, GPNMB, and SPP1 and have shown that these states are dynamically remodeled during steatohepatitis and fibrosis [30–35,38]. The present data therefore support a strong myeloid component in the bulk signature and provide a cellular context for genes that showed pronounced differences between the PC1-derived expression states. Individual signature genes also displayed distinct distributions across immune and stromal populations, offering candidates for future studies of immune-metabolic interactions in lipid-associated liver injury.

Independent dietary and pharmacologic cohorts supported cross-model responsiveness of the signature, an important consideration when assessing the reproducibility of a molecular signature beyond its discovery dataset [21,26]. In GSE287727, HFCF feeding increased the signature in both WT and HuRKO backgrounds, with stronger overall induction in HuRKO mice. Previous work has independently shown that hepatocyte HuR modulates hepatic lipid homeostasis during high-fat feeding [40]. One WT-HFCF animal showed a Low-signature-like pattern resembling G1, suggesting that attenuated individual transcriptional responses can also occur in an independent dietary model; the sample was retained in all formal analyses. In GSE235797, Ginkgetin produced a coordinated decrease in the signature, in agreement with the original study reporting improvement of NASH-associated molecular and cellular features after Ginkgetin treatment [41]. Because the Ginkgetin and Amlexanox cohorts differ in experimental design and measurement context, the magnitude of suppression should not be interpreted as a head-to-head comparison of drug efficacy.

Human genetic analyses further prioritized TAGLN2 as a translational candidate. Integrating cis-eQTLs with disease GWAS through Mendelian randomization and colocalization can help prioritize gene-trait relationships while addressing some limitations of conventional observational association, although these approaches remain dependent on instrumental-variable assumptions and the underlying regulatory resources [24,42–46]. Whole-blood TAGLN2 was the only tested gene-tissue signal that remained significant after FDR correction in the MR screen, and the TAGLN2-CHD region showed strong colocalization evidence. The other signature genes showed weaker or less consistent evidence across the source NAFLD GWAS. TAGLN2 has established roles in immune-cell actin organization and activation, and recent work has linked it to lipid-dependent immune-cell metabolism [47–49]. Notably, an independent proteome-wide MR and colocalization study also prioritized TAGLN2 among circulating proteins with genetic evidence for coronary artery disease [50]. Together, these findings support further investigation of TAGLN2 in cardiometabolic disease, while the MR and colocalization results should be interpreted as genetic evidence consistent with a causal relationship rather than definitive proof of direct gene-level causality.

This study has several limitations. All analyses were based on publicly available transcriptomic datasets, and direct phenotypic measurements were not available to determine whether the G1 Low-signature state reflects biological resistance to model induction or incomplete model establishment. The molecular regulators responsible for the PC1-defined heterogeneity also remain to be experimentally characterized. In addition, the single-cell dataset was enriched for non-parenchymal cells, and the human translational evidence was based on genetic inference rather than prospective clinical validation. Future work should integrate individual-level metabolic and histologic phenotypes, experimentally test candidate regulatory mechanisms, and evaluate the 20-gene signature and TAGLN2 in independent clinical cohorts.

## Conclusion

In summary, conventional treatment allocation did not fully capture the dominant hepatic transcriptional heterogeneity present in GSE338111. PC1-guided, treatment-independent stratification resolved three reproducible expression states and enabled derivation of a myeloid-associated 20-gene signature. The signature showed reproducible responses in independent dietary and pharmacologic cohorts and was predominantly localized to hepatic myeloid populations in single-cell data. Human genetic analyses identified TAGLN2 as the signature component with the strongest evidence linking the panel to coronary heart disease. These findings support PC1-guided stratification as a useful complementary strategy for transcriptomic datasets with substantial within-group heterogeneity and provide a focused molecular framework for future mechanistic and translational studies.

---

**Supplementary Figure S1.**
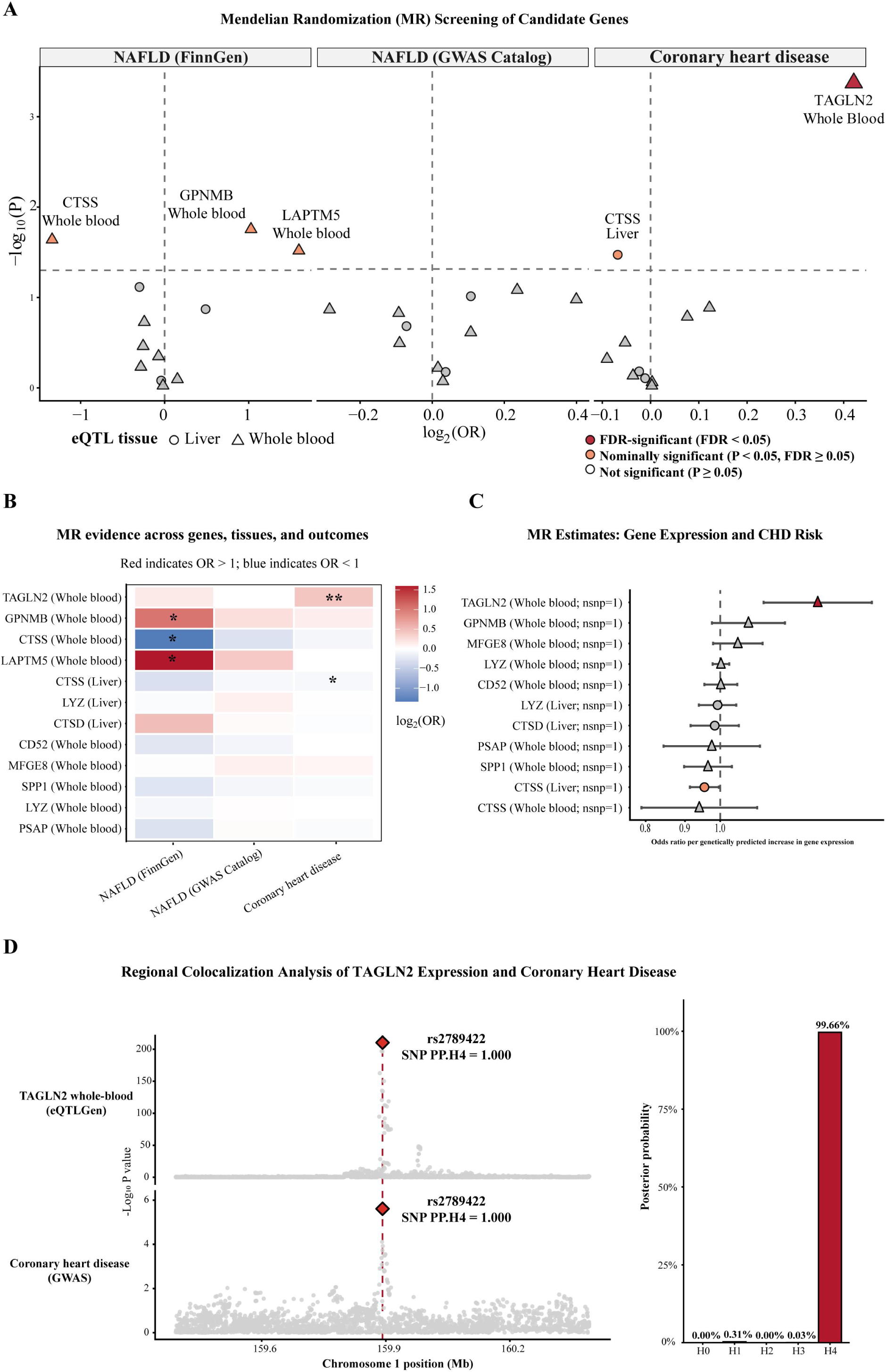
Original-treatment differential expression, transcriptome quality control, sensitivity to sex adjustment, and Pcsk9-related transcriptional assessment in GSE338111. (A) Volcano plot showing differential gene expression between Amlexanox- and DMSO-treated mice under the original treatment grouping. Differential expression was assessed using DESeq2 with sex included as a covariate (∼ sex + treatment). Genes with FDR < 0.05 and |log2FC| ≥ 1 were defined as stringent differentially expressed genes; 21 genes met these criteria. Genes belonging to the 20-gene signature derived from the PC1-guided analysis are highlighted. Dashed vertical lines indicate log2 fold changes of -1 and +1, and the horizontal dashed line indicates FDR = 0.05. (B) Sample-level quality-control metrics, including deposited count-like library size, number of detected genes, and the distribution of VST-transformed expression values. Samples are ordered by PC1-derived group. (C) Pairwise Spearman correlation matrices calculated from transcriptome-wide VST expression values. The same correlation matrix is shown with samples ordered by the original treatment assignment (left) and by the PC1-derived groups (right). (D) Concordance of genome-wide DESeq2 log2 fold-change estimates obtained from group-only and sex-adjusted models for G2 versus G1 and G3 versus G2. The 20-gene signature is highlighted, and Spearman and Pearson correlation coefficients are shown for each contrast. (E) Pcsk9 expression across the PC1-derived groups, shown as log2(DESeq2-normalized count + 1) for each mouse. Point color indicates PC1-derived group and point shape indicates sex, as shown in the figure. (F) Signed FXR/bile-acid and lipogenesis program scores across the PC1-derived groups. Individual mice are shown together with group means and SEM. (G) Heatmap of independent PCSK9/Paigen-related transcriptional programs after exclusion of the 20 candidate genes. Values are gene-wise VST Z scores, with sample annotations for sex, original treatment, and PC1-derived group.

